# Understanding multivariate brain activity: evaluating the effect of voxelwise noise correlations on population codes in functional magnetic resonance imaging

**DOI:** 10.1101/592618

**Authors:** Ru-Yuan Zhang, Xue-Xin Wei, Kendrick Kay

## Abstract

Previous studies have shown that neurons exhibit trial-by-trial correlated activity and that such noise correlations (NCs) greatly impact the accuracy of population codes. Meanwhile, multivariate pattern analysis (MVPA) has become a mainstream approach in functional magnetic resonance imaging (fMRI), but it remains unclear how NCs between voxels influence MVPA performance. Here, we tackle this issue by combining voxel-encoding modeling and MVPA. We focus on a well-established form of NC, tuning-compatible noise correlation (TCNC), whose sign and magnitude are systematically related to the tuning similarity between two units. We first replicate the classical finding that TCNCs impair population codes in a standard neuronal population. We then extend our analysis to fMRI data, and show that voxelwise TCNCs do not impair and can even improve MVPA performance when TCNCs are strong or the number of voxels is large. We also confirm these results using standard information-theoretic analyses in computational neuroscience. Further computational analyses demonstrate that the discrepancy between the effect of TCNCs in neuronal and voxel populations can be explained by tuning heterogeneity and pool sizes. Our results provide a theoretical foundation to understand the effect of correlated activity on population codes in macroscopic fMRI data. Our results also suggest that future fMRI research could benefit from a closer examination of the correlational structure of multivariate responses, which is not directly revealed by conventional MVPA approaches.

## INTRODUCTION

Understanding how neural populations encode information and guide behavior is a central question in cognitive neuroscience. In a neuronal population, many units exhibit correlated activity, and this likely reflects an important feature of information coding in the brain. In computational neuroscience, researchers have investigated the relationship between *signal correlation* (SC), referring to the similarity between the tuning functions of two neurons, and *noise correlation* (NC), referring to the correlation between two neurons’ trial-by-trial responses evoked by repetitive presentations of the same stimulus (Averbeck et al., 2006; Cohen & Kohn, 2011; Kohn et al., 2016).

Prior studies in neurophysiology have discovered that neurons that share similar tuning functions (i.e., a positive SC) also tend to have a weak positive NC, a pervasive phenomenon has been across several brain regions (Averbeck & Lee, 2003; Constantinidis & Goldman-Rakic, 2002; Ecker et al., 2010; Gutnisky & Dragoi, 2008; Huang & Lisberger, 2009; Jermakowicz et al., 2009; Lee et al., 1998; Smith & Kohn, 2008). In this paper, we denote this type of NC as *tuning-compatible noise correlation* (TCNC) because the sign and the magnitude of the NC are systematically related to the SC between a pair of neurons. A bulk of theoretical and empirical work has shown that NCs have a substantial impact on population codes. For example, the seminal study by Zohary et al. (1994) demonstrated that TCNCs limit the amount of information in a neural population as the noise is shared by neurons and cannot be simply averaged away. Later on, researchers realized that this detrimental effect of TCNC is mediated by other factors, such as the form of NC, heterogeneity of tuning functions and its relevance to behavior (Haefner et al., 2013).

The study of NCs in the brain has been historically impeded by technical barriers to measuring simultaneously the activity of many neurons in neurophysiological experiments. In contrast, functional magnetic resonance imaging (fMRI) naturally measures the activity of many neural populations throughout the entire brain. Imaging scientists often use multivariate pattern analysis (MVPA) to assess the accuracy of population codes (Haxby et al., 2014; Tong & Pratte, 2012). However, above-chance decoding performance in MVPA does not specify the detailed representational structure underlying multivariate voxel responses. For example, Fig. 1 illustrates a simple two-voxel scenario in multivariate decoding. The decoding accuracy in the original state (Fig. 1A) can be improved (e.g., by attention, learning) via either the further separation of mean responses (Fig. 1B) or the changes to the covariance geometry (Fig. 1C). This example highlights the impact of the shape of the response distribution on population codes and these effects cannot be easily disentangled by the conventional MVPA approach (Naselaris & Kay, 2015).

**Figure 1.**
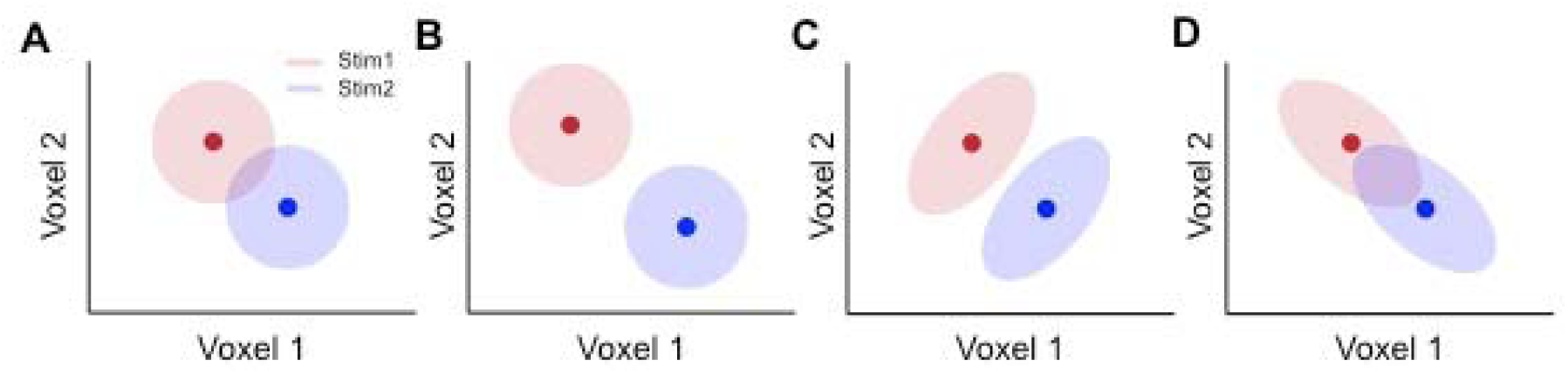
A two-voxel scenario in MVPA. The pool consists of two responsive voxels and two color disks represent the trial-by-trial response distributions evoked by two different stimuli. Panel A illustrates the original state of the population responses. Decoding performance can be improved via either a bigger separation of the mean population response (panel B) or changes in the covariance structure (panel C). Representational structures in panels B and C indicate improved population codes but have distinct underlying mechanisms. Panel D illustrates that certain covariance changes can worsen decoding.

The magnitude and the structure of NCs in fMRI data still remain largely unknown. It has been shown that NCs influence MVPA accuracy and that certain types of classifiers can compensate for NCs (Cox & Savoy, 2003). But the precise nature of NCs has not yet been thoroughly characterized. There have been a few recent investigations of NCs. A study by Ryu and Lee (2018) evaluated the impact of three factors—retinotopic distance, cortical distance, and tuning similarity—on voxelwise NCs in early visual cortex, and found that tuning similarity is the major determinant for voxelwise NCs. Furthermore, van Bergen and Jehee (2018) systematically evaluated voxelwise NCs in human V1 to V3 and showed that the magnitude of NCs monotonically increases as tuning similarity increases. They also found that voxelwise NCs in the striate cortex are in general higher than NCs in the extrastriate cortex. Furthermore, one recent study found that a multivariate classifier can exploit voxelwise NCs to decode population information (Bejjanki et al., 2017). These results provide specific evidence supporting the existence of voxelwise TCNC, and suggest that a deeper understanding of how NC manifests in fMRI data is critical for studying probabilistic neural computation using multivariate fMRI data (Ma & Jazayeri, 2014; van Bergen & Jehee, 2018).

In the present study, we combine MVPA and the voxel-encoding modeling approach to assess how the magnitude and form of NCs impact population codes in fMRI data. Similar to prior theoretical work in neurophysiology, we aim to derive the theoretical bound of the effects of voxelwise NCs on population codes in multivariate voxel responses. We assess the accuracy of population codes by MVPA and information-theoretic analyses. The voxel-encoding model used in this study allows us to systematically manipulate response parameters (i.e., voxel tuning) so as to examine NCs in different scenarios (Naselaris et al., 2011). We first assess the quantitative relationship between decoding accuracy and the strength of NCs. We then directly calculate the amount of information as a function of NCs in a voxel population. Both methods demonstrate that the accuracy of population codes in fMRI data follows a U-shaped function as the strength of TCNC increases. Notably, all these analyses in voxel populations are compared against classical findings in neuronal populations. We show that the effects of NCs on population codes are strongly mediated by tuning heterogeneity in both neuronal and voxel populations.

## RESULTS

### 1. Effects of noise correlation in neuronal and voxel populations on both stimulus estimation and classification

In the first part, we will show the effects of noise correlation on population codes in both a neuronal and a voxel population. We simulated multivariate responses with three and two forms of noise correlation in neuronal and voxel populations respectively. We performed two brain decoding tasks — a stimuli-classification task and a stimulus-estimation task. In the stimulus-classification task, a linear classifier was trained to categorize evoked population responses into one of two discrete orientation stimuli. In the stimulus-estimation task, a maximum likelihood estimator (MLE) was used to reconstruct the continuous orientation value based on the population response in a trial. Both tasks are two routinely used forms of MVPA in the literature (Haxby et al., 2014; Kamitani & Tong, 2005; Tong & Pratte, 2012).

#### 1.1 Tuning-compatible noise correlations between neurons impair decoding accuracy in multivariate neuronal responses

Before examining the effect of TCNC in a voxel population, we first attempted to replicate the classical findings in a standard neuronal population. In the simulation of neuronal population responses, all neurons shared the same tuning curve except that their preferred orientations were equally spaced in the continuous orientation space (Fig. 2A, also see Methods). We manipulated three types of NCs between neurons—angular-based tuning-compatible noise correlation (aTCNC), curve-based tuning-compatible noise correlation (cTCNC) and shuffled noise correlation (SFNC). The first one is also called ‘limited-range correlation’ in neurophysiological literature (Cohen & Kohn, 2011; Zohary et al., 1994). aTCNCs are based on the angular difference between the preferred orientations of two neurons (Fig. 3A). Specifically, we defined that the strength of NC between two neurons follows an exponential decay function (Eq. 2) of the absolute angular difference between their preferred orientations. This approach has often been used to establish population coding models (Bair et al., 2001; Ecker et al., 2011; Zohary et al., 1994). The second type, cTCNC, is based on the similarity between the tuning curves (i.e., SC) of two neurons (Fig. 3B). We defined that the sign and magnitude of cTCNCs are the same as and proportional to the SC between two neurons. This is consistent with empirical measurement in electrophysiology (Bair et al., 2001; Ecker et al., 2011; Zohary et al., 1994). Note that both aTCNC and cTCNC are related to the tuning similarity between two neurons since the larger angular difference between the two neurons’ preferred orientations, the less their tuning curves are correlated. SFNCs served as a control condition and were generated by randomly shuffling the cTCNCs between neurons (Fig. 3C, see Methods) such that they had no relationship with the tuning properties of neurons.

**Figure 2.**
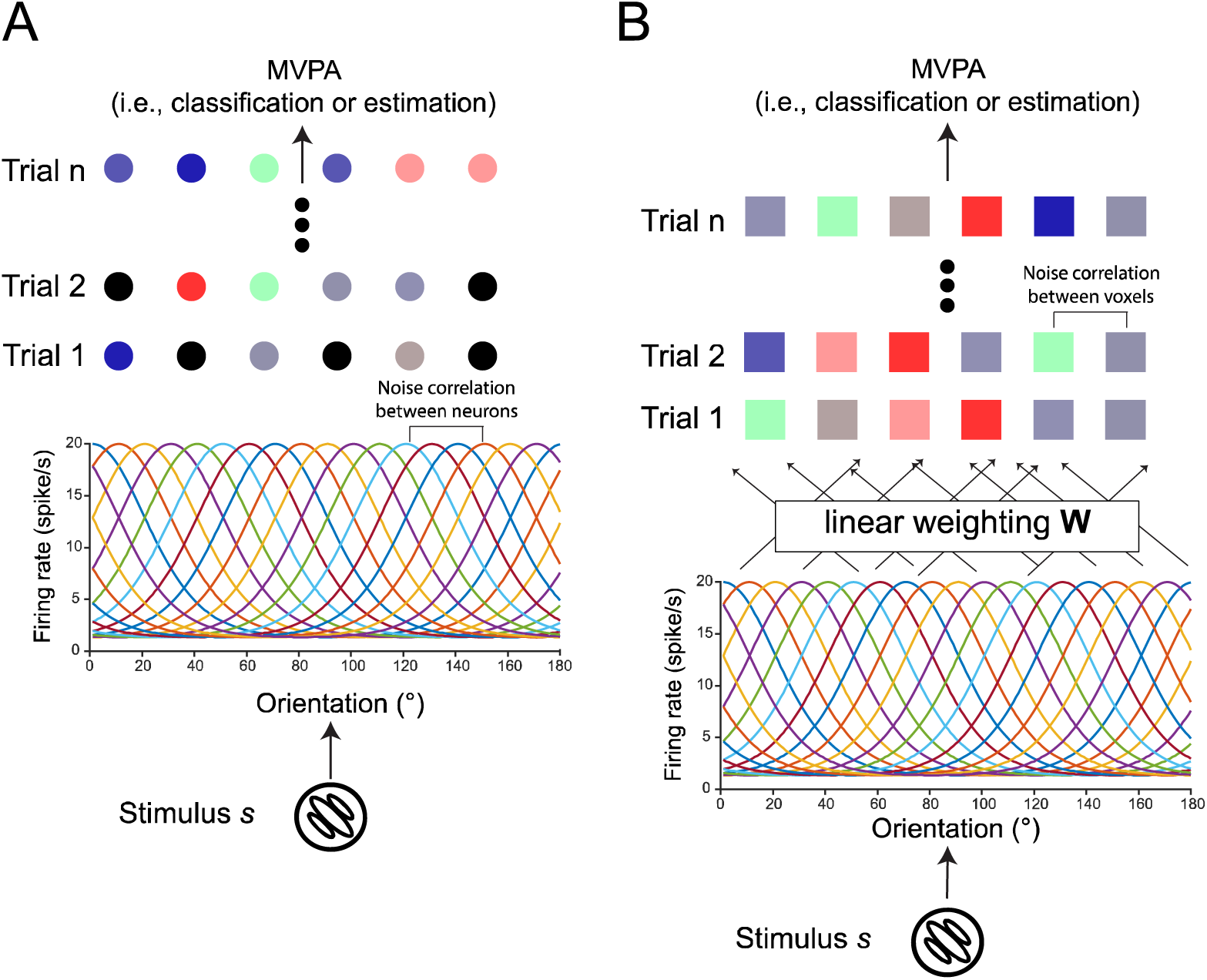
Neuron- and voxel-encoding models. The neuron-encoding model (panel A) proposes a neuronal population with homogenous orientation-selective tuning curves. Each neuron has Poisson-like response variance and the noise correlation between two neurons can be specified with different structures and strength (see Methods). The voxel-encoding model proposes a similar neuronal population and the response of a single voxel is the linear combination of the responses of multiple neurons. The noise correlation between two voxels can be specified using similar methods (see Methods). Note that voxelwise NCs can come from the response variability at both neuronal and voxel levels (see Fig. 6). Using the neural- and the voxel-encoding model, we can generate many trials of neuronal and voxel population responses and perform conventional MVPA on the simulated data. The goal is to examine multivariate decoding performance as a function of the NC structure and strength between either neurons or voxels.

**Figure 3.**
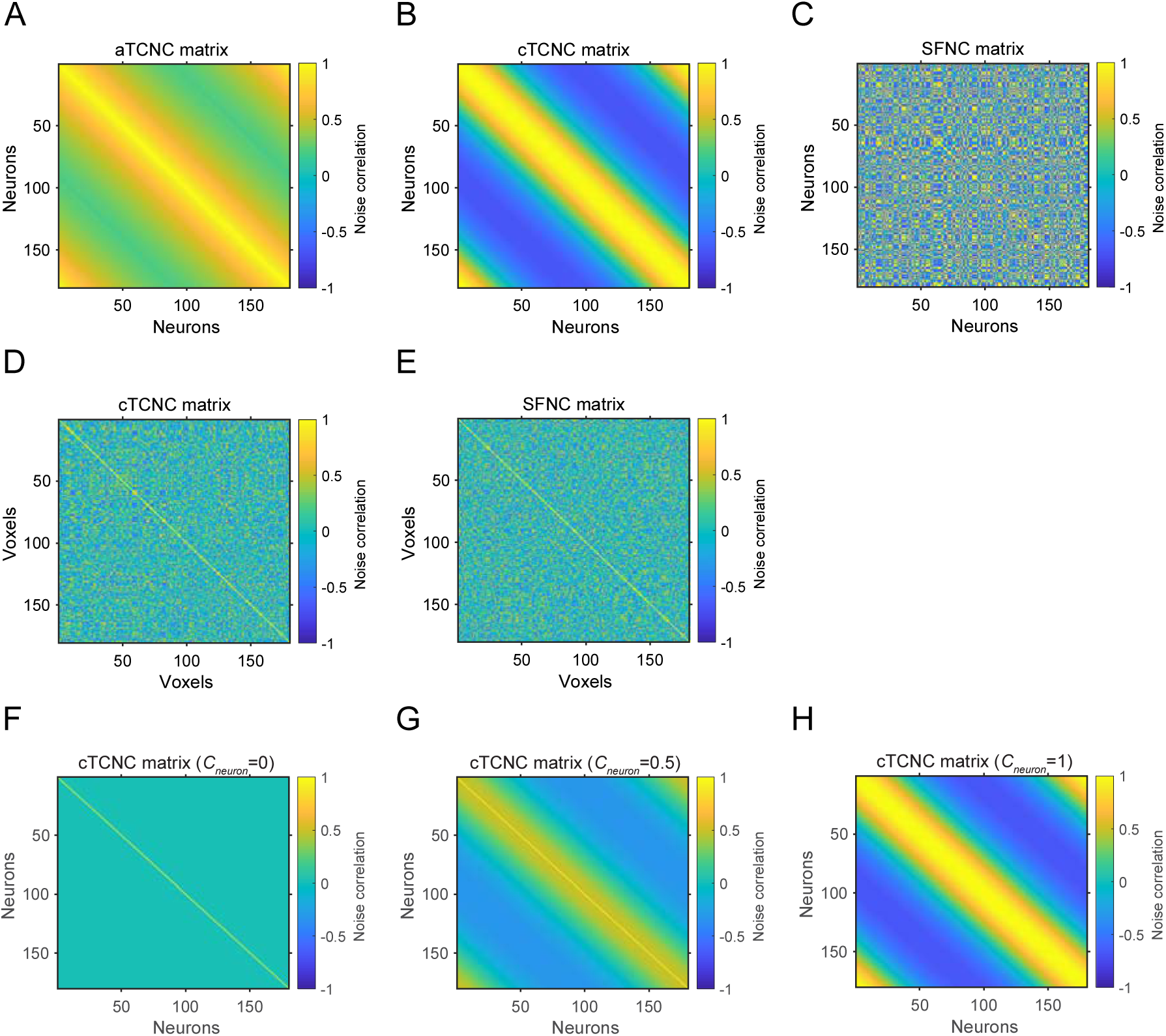
Example noise correlation matrices simulated in a neuronal (panels A-C) and a voxel population (D, E). In the neuronal population (180 neurons), the angular-based TCNC matrix, the curve-based TCNC matrix, and the SFNC matrix are illustrated from left to right. Neurons are sorted according to their preferred orientation from 1 to 180°. In the voxel population (180 voxels), the curve-based TCNC matrix and the SFNC matrix are illustrated. Note that we do not sort the voxels according to their tuning preference. The NC coefficients (*c*_*neuron*_ or *c*_*vxs*_) are set to 1 in matrices from A-E. Panels F-H illustrate the cTCNC matrices with NC coefficient (*c*_*neuron*_) values 0, 0.5 and 1, respectively. Note that panels B and H are identical.

We manipulated two variables of the neuronal population—the pool size (i.e., the number of neurons) and the strength of NCs between neurons (i.e., *c*_*nc*_, see Fig. 3F-H). For every combination of pool sizes and NC strength levels, we simulated population responses in many trials and performed the MVPA decoding (i.e., classification and estimation) on the simulated population responses.

Results indeed replicated the findings from previous theoretical work (Zohary et al., 1994). aTCNCs and cTCNCs impaired decoding performance in both tasks: the classification accuracy (Fig. 4A-B) and the efficiency of the MLE (Fig. 4D-E) declined as the strength of aTCNCs and cTCNCs increased. Decoding performance always rose as the strength of SFNCs increased. This result is similar to finding in (Wilke & Eurich, 2002). We will explain this phenomenon in the later sections.

**Figure 4.**
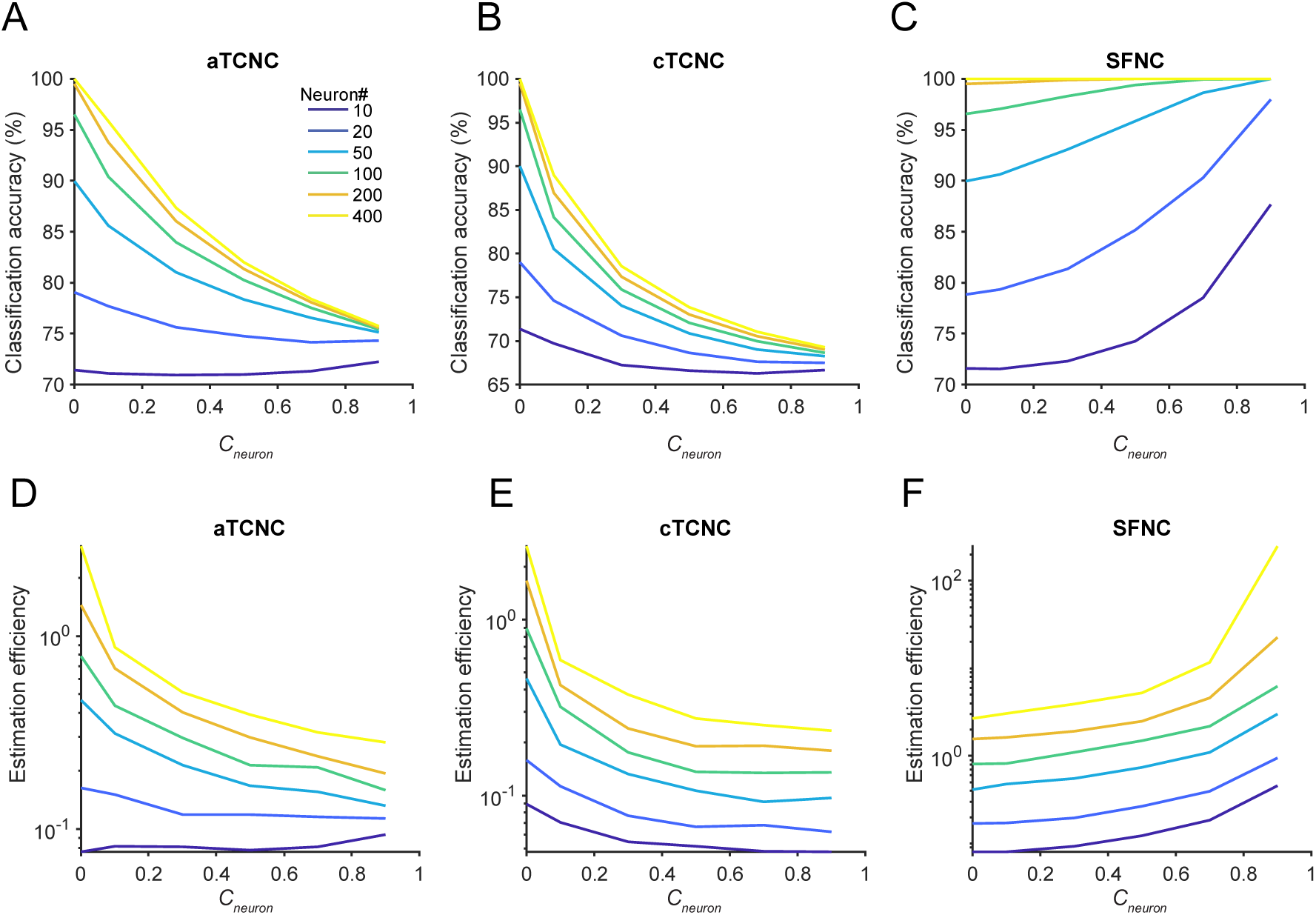
TCNCs impair population codes in a neuronal population. The multivariate classification accuracy (panels A-C) and maximum likelihood estimation efficiency (panels D-F) are depicted as a function of the magnitude of the aTCNC (panels A, D), TCNC (panels B, E) and the SFNC (panels C, F). Both classification accuracy and estimation efficiency decline as the strength of aTCNC and cTCNC increases. Conversely, increasing the strength of SFNC improves decoding accuracy.

#### 1.2 Decoding accuracy as U-shaped functions of tuning-compatible noise correlations in multivariate voxel responses

We next turned to examine the impact of NC on population codes in fMRI data. We simulated responses of a voxel population using a voxel-encoding model (Fig. 2B) and attempted to perform the classification and estimation tasks.

We again manipulated two types of NCs—cTCNC (Fig. 3D) and SFNC (Fig. 3E). The cTCNCs here are similar to above except that they are between voxels rather than neurons. Similarly, cTCNCs here are defined with respect to the similarity of their orientation tuning curves. SFNCs were also generated by randomly shuffling the cTCNCs between voxels (see Methods). Note that we cannot parametrically derive aTCNCs for voxels as we did for neurons since unlike unimodal orientation tuning curves of cortical neurons, orientation tuning curves of voxels might be irregular (i.e., multimodal) due to the mixing of multiple neural populations in a voxel’s activity (see Eq. 9). We will return to this point in a later section.

Surprisingly, we found that the decoding performance exhibited U-shaped functions of the increasing amount of cTCNCs: both classification accuracy (Fig. 5A) and estimation efficiency (Fig. 5C) first declined and then rose in both tasks. This is puzzling since the predominant view in neurophysiology regards cTCNCs as detrimental but here we demonstrated that cTCNCs improve population codes. SFNCs in general improved the decoding accuracy, similar to the effect observed in a neuronal population.

**Figure 5.**
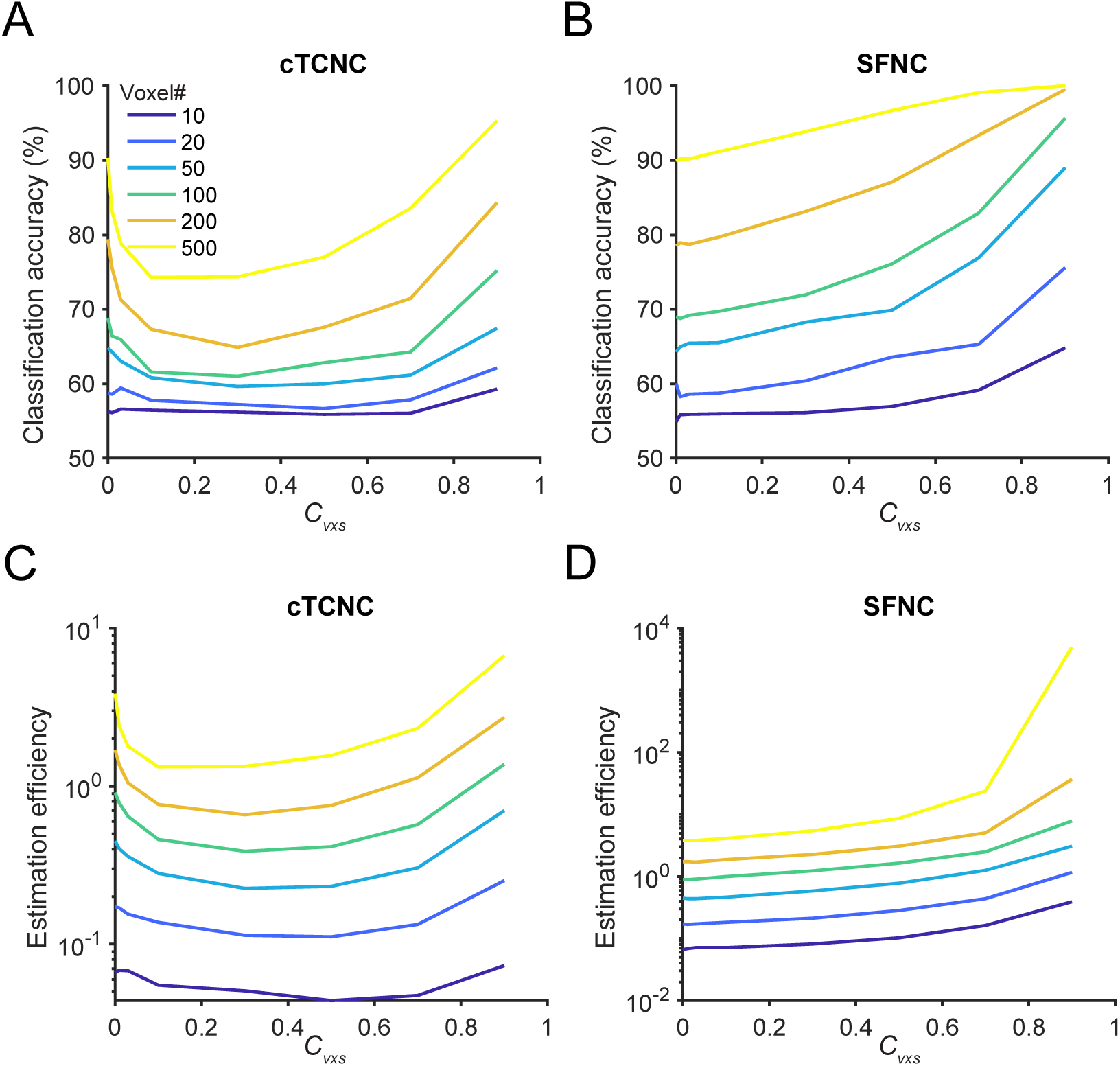
Decoding accuracy as U-shaped functions of cTCNCs in a voxel population. The multivariate classification accuracy (panels A, B) and estimation efficiency (panels C, D) are depicted as a function of the magnitude of cTCNCs (panels A, C) and SFNCs (panels B, D). Decoding accuracy exhibits U-shaped functions as cTCNCs increase. Similar to a neuronal population, SFNCs always improve decoding accuracy.

#### 1.3 Simultaneously varying neuronal and voxelwise noise correlations

In empirical fMRI studies, we can only measure voxelwise NCs but the sources of these NCs are unclear. One important source might be neuronal NCs because neuronal NCs could propagate to the voxel level if voxel responses are believed to be the aggregation of neuronal responses. However, fMRI data might also involve other MRI-specific noises (e.g., hemodynamic fluctuations, thermal noise, head motion, etc.). It is thus reasonable to assume that voxel-level NCs reflect the combinations of neuronal and other voxel-level NCs. Systematically disentangling these factors would be a useful direction for future experimental studies, but here we can at least derive some theoretical expectations using our analytical framework. In previous analyses, we only manipulated either the NCs between neurons or the NCs between voxels. We next manipulate both neuronal and voxelwise cTCNCs in the voxel-encoding model.

We repeated the classification and the estimation tasks on a voxel population (see Methods for details). Results showed that increasing neuronal-level cTCNCs had a small impact on classification accuracy and the change in the classification accuracy values was primarily determined by voxel-level cTCNCs. This is because we attempted to decode two stimuli (*s*_*1*_ = 80°, *s*_*2*_ = 100°) based on simulated fMRI responses. But this is a very easy task if we classify the two stimuli directly from neuronal responses (i.e., reach 100% correct ceiling, also see Methods). Thus, classification accuracy here is primarily bottlenecked by the noise at the voxel level not the neural level. Note that these results are contingent on the noise structure and strength assumed at both processing stages.

In the stimulus-estimation task, neuronal cTCNCs dampened estimation efficiency and voxelwise cTCNCs impact estimation efficiency as U-shaped functions. Both results are consistent with the previous results when two levels of NCs were manipulated independently.

### 2. Results of information-theoretic analyses explain the effects of noise correlation on population codes

In the second part, we will show how to use information-theoretic analyses to support the simulation results above. Especially, we want to highlight the unit tuning heterogeneity as a mediator for the effect of NCs in a population. Unit tuning heterogeneity also acts as the key factor to explain the differential effects of cTCNC in neuronal and voxel populations.

#### 2.1 Amount of information echoes decoding accuracy in population codes

Above analyses focused on assessing the population codes from the decoding perspective (i.e., MVPA), the approach that almost all previous fMRI decoding studies used. Here, we propose an alternative approach—directly calculate the amount of Fisher information for the stimulus-estimation task and linear discriminability for the stimulus-classification task. They have been used as the standard metric for information coding in computational neuroscience (Brunel & Nadal, 1998; Seung & Sompolinsky, 1993). For the estimation task, Fisher information indicates the minimal amount of variance that any unbiased decoder can possibly achieve. For the classification task, linear discriminability measures the magnitude of separation of two multivariate response distributions. It is also called a variant of linear Fisher information for a classification task (Kohn et al., 2016). For simplicity, we termed both metrics as “information” as they both indicate the accuracy of population codes with respect to the two tasks.

The analysis of information has three major advantages over the conventional MVPA approach. First, in theory two approaches might lead to consistent results as more information in a population usually leads to a higher decoding accuracy. But their relationship is nonlinear. Classification accuracy can reach the floor (e.g., 50% for binary classification) and ceiling (i.e., 100%) but the amount of information has a relatively broad range thus more sensitive to population codes. For example, as we will show, information in a standard neuronal population saturates as a function of pool size given the presence of aTCNCs and cTCNCs but not SFNCs (Fig. 7F-H). Information in a voxel population keeps increasing as the pool size increases (Fig. 7F-J). these conclusions cannot be easily derived from decoding analyses per se. Second, unless optimal Bayesian decoders are used, decoding performance is not guaranteed to produce the same results as analysis of information. Researchers mostly used some machine learning methods, such as support vector machine, logistic regression, linear regression. It still remains unclear whether these decoders are statistically optimal. The decoding results above might due to the particular decoders we use. In contrast, the assessment of information is not related to the assumptions or efficacy of any particular decoder. Third, most decoding methods so far employed discriminative modeling approach. Calculation of information here takes into account data generative processes. As such, calculation of information should be a more principled way to assess the accuracy of population codes.

We calculated the amount of information in both the neuronal and the voxel populations (see Methods) as functions of pool size and NC strength. Results largely mirrored the previous decoding results. In the neuronal population, we replicated the key signatures of detrimental effects of TCNCs: the amount of information saturated as the pool size increased given the presence of aTCNCs (Fig. 6F) and cTCNCs (Fig. 6G) but not SFNCs (Fig. 6H). Also, the amount of information declined as the magnitude of aTCNCs (Fig. 6A) and cTCNCs increased (Fig. 6B). This pattern was reversed as the magnitude of SFNCs increased (Fig. 6C). In the voxel population, the amount of information always increased as the pool size expanded in both cTCNCs (Fig. 6I) and SFNCs (Fig. 6J) conditions. Similar to the decoding results, the amount of information exhibited U-shaped functions as the magnitude of cTCNCs (Fig. 6D) increased and always grew as the magnitude of SFNCs (Fig. 6E) increased.

**Figure 6.**
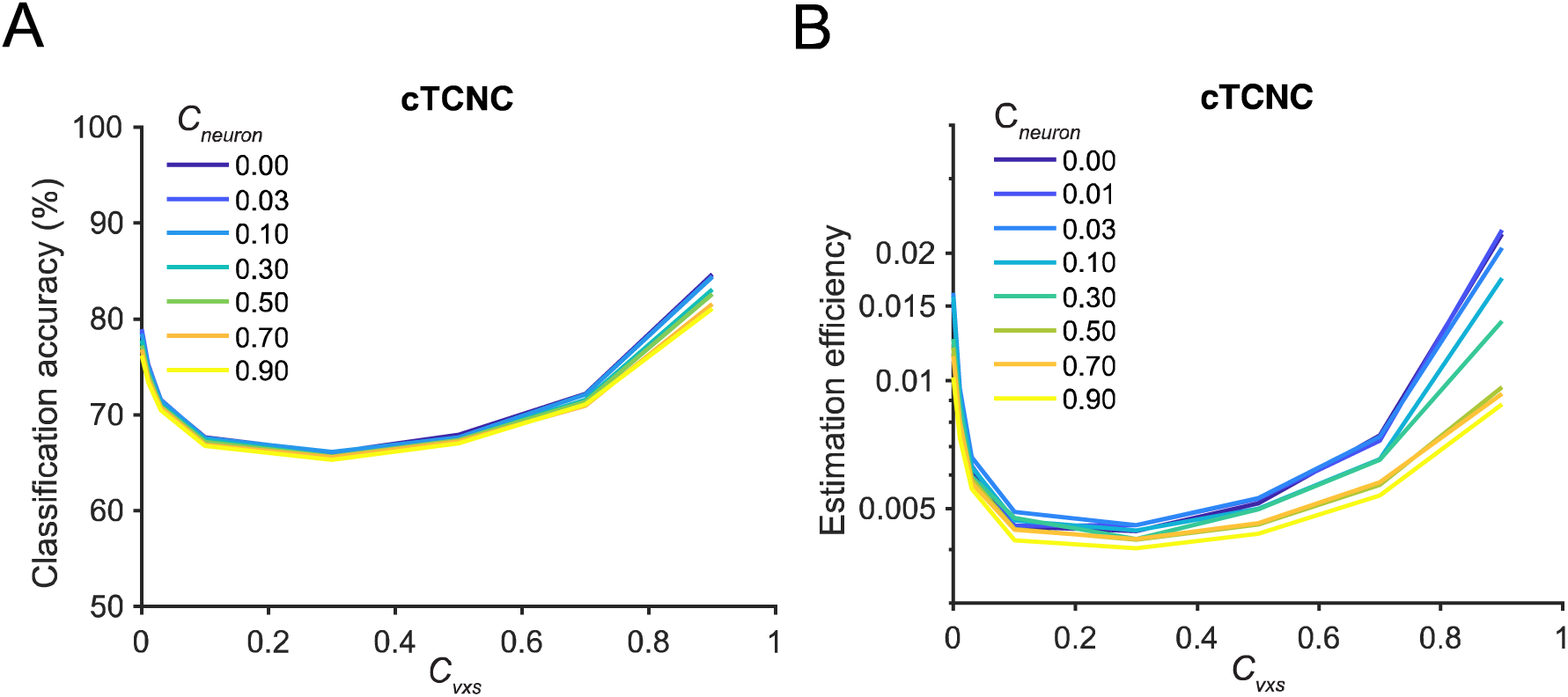
The impacts of neuronal and voxelwise cTCNCs on stimulus classification (A) and estimation (B). In both tasks, decoding performance exhibits U-shaped functions of the strength of voxelwise cTCNCs (i.e., *c*_*vxs*_). Neuronal cTCNCs (i.e., *c*_*neuron*_) have small impacts on classification accuracy, because the voxel-level noise primarily limits information. Neuronal cTCNCs have a more prominent detrimental effect in the estimation task. These results are consistent with the results when two levels of cTCNCs are manipulated independently.

#### 2.2 Voxel tuning heterogeneity and pool sizes explain the effect of tuning-compatible noise correlations on population codes

Why do TCNCs manifest differently in neuronal and voxel populations? We reason that the neuron-to-voxel transformation (i.e., linear weighting **W**) might be the key factor that alters the effect of TCNCs. Unlike the homogeneous neuronal tuning curves (i.e., same width, amplitude, and baseline, and only preferred orientations vary), voxel tuning curves might be heterogeneous or have diverse tuning widths and amplitudes (Fig. 8C). This is due to the uncertain distribution (i.e., the weighting matrix **W** in Eq. 9) of orientation-selective neurons within a voxel. Even though individual neurons follow a uniform bell-shape tuning property, the aggregation of them can produce tuning functions with diverse forms. Because of the tuning heterogeneity, TCNCs do not limit information anymore. The effect of tuning heterogeneity has been studied in some previous theoretical work (Ecker et al., 2011; Shamir & Sompolinsky, 2006; Wilke & Eurich, 2002) (see more details in discussion).

To further substantiate the interaction effect between tuning heterogeneity and NCs on population codes, we performed two additional analyses. First, in the voxel population (Fig, 7A), we normalized the information when cTCNCs are present (i.e., *c*_*vxs*_ > 0) by the information when cTCNCs are absent (i.e., *c*_*vxs*_ = 0). We found a similar pattern as in a heterogeneous neuronal population (Fig. 8B) as suggested by a previous study (Ecker et al., 2011). Importantly, the relative benefits of TCNCs are all U-shaped functions. Such beneficial effects are more pronounced in a large population (i.e., the lowest point of the function shifts to left as the pool size increases in Fig. 8A).

**Figure 7.**
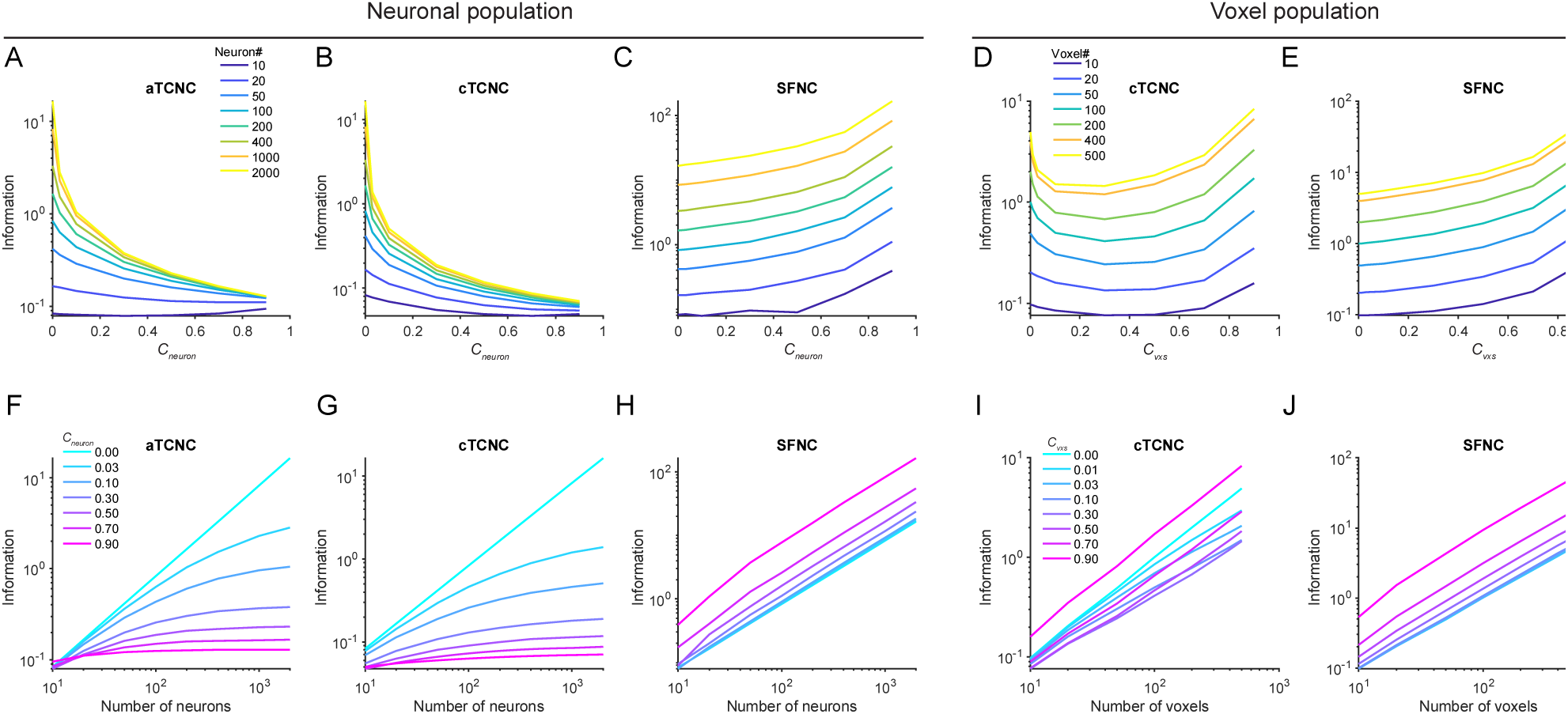
Amount of information in neuronal and voxel populations with diverse forms and strength of NCs. The upper and the bottom rows depict the amount of information as a function of increasing strength of NCs and the increasing number of units in the population, respectively. Panels A-C and F-H illustrate the amount of information in a neuronal population and correspond to Fig. 4. Panels D-E and I-J illustrate the amount of information in a voxel population and correspond to Fig. 5. Note that here we only illustrate the information in a stimulus-estimation task. We consider three types of NC—aTCNC (panels A, F), cTCNC (panels B, G) and SFNCs (panels C, H) in the neuronal population, as already shown in Fig. 4. Similar treatments are performed for the voxel population, as shown in Fig. 5. The calculation of information largely mirrors the decoding results shown in Fig. 4 and Fig. 5. Critically, the amount of information in the voxel population exhibits U-shaped functions of increasing strength of cTCNCs (panel D) and cTCNCs do not limit information as the number of voxels increases (panel I). These results clearly differ from the effects of aTCNCs (panels A, F) and cTCNCs (panels B, G) in a neuronal population.

**Figure 8.**
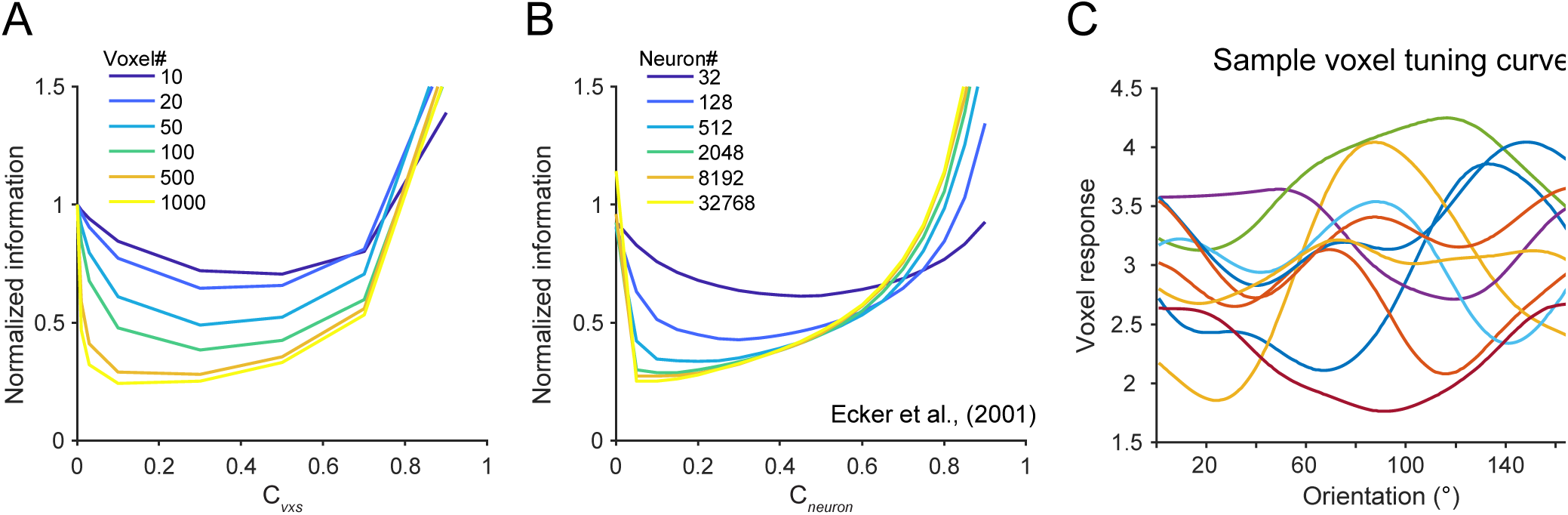
Tuning heterogeneity might be the key mediator for the effect of TCNC in a voxel population. In panels A, the y-axis indicates the amount of information when NCs are present normalized by the amount of information when NCs are 0. Normalized information in our simulation is directly reminiscent of the information on a neuronal population with heterogeneity tuning curves (panel B, reproduced from (Ecker et al., 2011)). This result implies that tuning heterogeneity mediates the effect of TCNC in both types of data. Panel C illustrates some sample tuning curves of the simulated voxels. Due to the uncertain neuron-to-voxel connections (i.e., linear weight matrix **W**), the endowed voxel tuning curves also exhibit irregular forms.

In the second analysis, we manipulated the degree of voxel tuning heterogeneity and the strength of cTCNCs in the voxel population. The amount of information was calculated as a function of these two variables (Fig. 9A&B). Results showed that the amount of information follows U-shaped functions if voxel tuning is highly heterogeneous (i.e., *c*_*homo*_=0.03 in Fig. 9). However, as the voxel tuning becomes progressively homogeneous (i.e., *c*_*homo*_ increases to 1), cTCNCs become more and more detrimental for information coding, which is consistent with the results obtained in a standard neuronal population (Fig. 7B). Taken together, we demonstrated that unit tuning heterogeneity is the key factor that mediates the contribution of cTCNCs in both neuronal and voxel populations.

**Figure 9.**
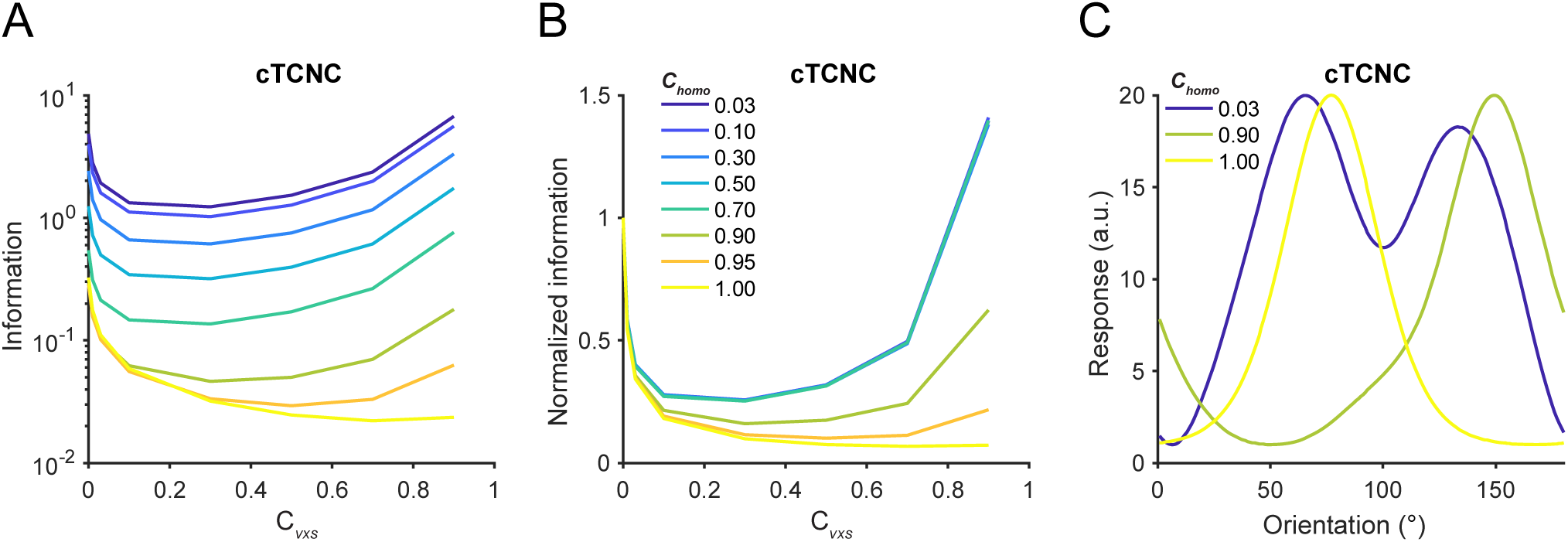
Interaction between cTCNC and tuning heterogeneity on population codes. 500 voxels were simulated (see Methods). A larger value of *c*_*homo*_ indicates more homogeneous voxel tuning curves. Note that the simulated voxel tuning curves are identical to neuronal tuning curves when *c*_*homo*_ =1. Panels A and B illustrate the raw and normalized amount of information as a function of cTCNC under different tuning heterogeneity levels. The raw information is normalized to the condition when *c*_*vxs*_=0 (panel B). As voxel tuning homogeneity increases, the shape of the functions changes from U-shaped to monotonically decreasing. Panel C illustrates sample voxel tuning curves with different heterogeneity levels.

#### 2.3 Winner-take-all principle as an intuitive explanation for the effect of shuffled noise correlations on population codes

Besides the beneficial effect of TCNCs, we turn to another interesting finding—SFNCs improve population codes in both neuronal and voxel populations. At first glance, this seems surprising since it suggests that decoding accuracy can be improved by randomly injecting some NCs between voxels. Here we want to highlight an intuitive explanation—a winner-take-all principle enhances decoding accuracy in the conventional multivariate analysis.

We simulated a simple three-voxel scenario for a classification task to illustrate this effect (Fig. 10). The NC between voxels X and Y improves classification (Fig. 10A), while the NC between voxels Y and Z impairs classification (Fig. 10B). The correlation between X and Y, and the correlation between Y and Z are identical in magnitude but with opposite signs. However, when all three voxels are aggregated, the contributions of two opposite NCs do not cancel out each other and the overall decoding performance is still improved by the positive NC between X and Y, regardless of the negative NC between Y and Z. Importantly, classification accuracy on X, Y, and Z with NCs is higher than the scenario in which there are no noise correlations. These results demonstrate that, as long as there exists at least one pair of “good” voxels whose NC is beneficial, a linear classifier can utilize the NC to achieve a good decoding performance. In other words, the most informative units determine the decoding accuracy that a decoder that can possibly achieve (Fig. 10).

**Figure 10.**
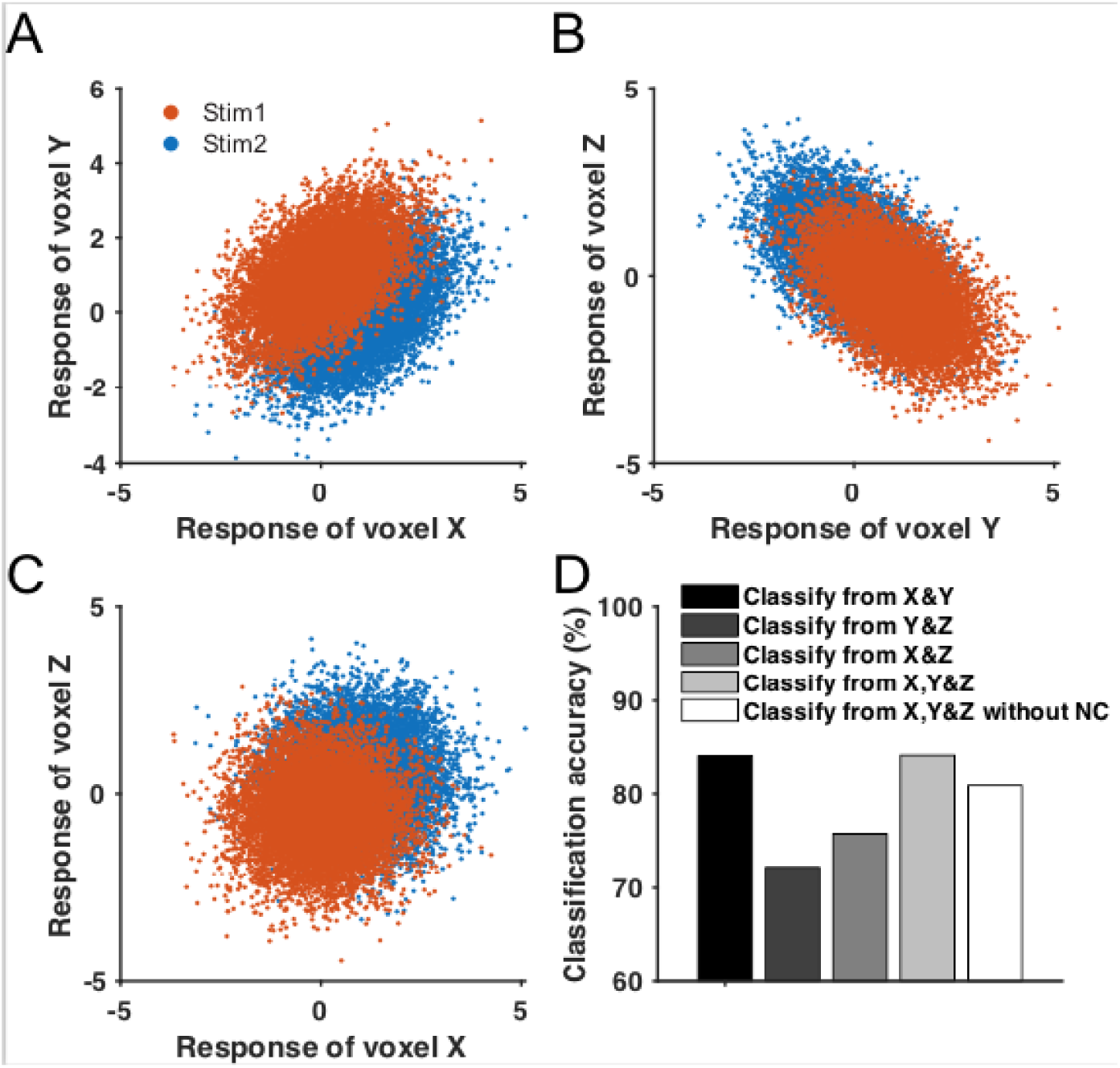
A three-voxel simulation illustrating the winner-take-all principle of multivariate decoding. Panel A illustrates the trial-by-trial responses of voxels X and Y towards two stimuli. The covariance structure of X and Y enhances classification accuracy. Similarly, panel B illustrates that the covariance structure of voxels Y and Z impairs classification. Voxels X and Z have no systematic NC (panel C). Panel D depicts the classification accuracy based on population responses of X and Y (panel A), Y and Z, X and Z, and all three units, respectively. We also include a situation where we set all NCs among three units to 0 and keep other settings the same. We add this condition because in most empirical scenarios we are interested in comparing a population code with and without NCs. The beneficial and detrimental effects of the covariance structures in panels A and B do not cancel each other if all three voxels are combined. The decoding performance of voxels X and Y determines the classification accuracy based on all three voxels (i.e., winner-take-all principle).

This winner-take-all principle provides an intuitive explanation for the effects of SFNCs on decoding. SFNCs are generated by randomly shuffling the TCNCs between voxels such that the NCs between some pairs of units might be beneficial whereas in other cases might be detrimental when considered individually. However, the overall decoding accuracy is related to the most beneficial NCs. As the pool size increases, it is more likely that a small portion of units can be assigned with NCs that greatly benefit decoding, resulting in an overall enhanced decoding accuracy on the entire population.

## DISCUSSION

Characterizing the effect of noise correlation on population codes has attracted much attention in the past years as it is related to several key topics (e.g., probabilistic computation, uncertainty) in neuroscience research (Kohn et al., 2016; Ma & Jazayeri, 2014). But the majority of relevant studies are confined to the field of neurophysiology. On the other hand, fMRI can measure many responsive units in the brain but most prior fMRI studies only employed MVPA to evaluate population codes. MVPA accuracy, however, is merely a coarse description of population codes and the precise quantitative relationship between the voxelwise NCs and population codes still remains unclear. Here, we conducted a series of theoretical analyses to systematically examine how NCs with different forms and strength influence MVPA accuracy and the amount of information in multivariate fMRI responses. We made three major observations: (1) decoding accuracy and the amount of information follow U-shaped functions of cTCNCs in a voxel population and this effect is mediated by voxel tuning heterogeneity and pool sizes; (2) even randomly injecting NCs between voxels improves population codes and this effect can be intuitively understood through a simple winner-take-all principle of decoding; (3) furthermore, the comparisons against a standard neuronal population demonstrate that the effect of NC in both neuronal and voxel populations can be understood within a unified computational framework related to tuning heterogeneity.

### Noise correlation in neural processing

The effects of NC on the capacity of the neural population code have been investigated in various studies over the past two decades (e.g., Abbott and Dayan, 1999; Sompolinsky et al., 2001; Shamir and Sompolinsky, 2006; Ecker et al., 2011), leading to somewhat mixed results. Early results in neurophysiology suggest that cTCNCs could be detrimental (Zohary et al., 1994), but later studies suggest that the results may be more complicated, depending on the detailed configurations of the neural codes. There are regimes where the cTCNCs could be beneficial (Wilke and Eurich (2002), Shamir and Sompolinsky, 2006; Ecker et al., 2011). Wilke and Eurich (2002) found that adding jitter to a uniform positive correlation structure (i.e., making the correlation more random) would benefit the code. They further provided an intuitive argument on why noise correlations that have no direct relationship to unit tuning might increase the coding capacity. Their results are consistent with our findings on the benefit of SFNCs.

The assumption of homogeneous tuning curves in early theoretical work is apparently not realistic because in the primate brain it has been known that the shape of tuning curves varies drastically across neurons. Such tuning heterogeneity removes TCNCs’ limitation on information. In a very large population, TCNCs even always benefit the population codes (Fig. 8B). This theoretical implication has been also corroborated by an empirical study on orientation decoding in primate V1 (Graf et al., 2011). Most importantly, the principle of tuning heterogeneity applies both neuronal and voxel populations.

Ecker et al. (2011) derived a mathematical foundation for the effects of tuning heterogeneity, pool size, and TCNC on population codes, built up the earlier work by Sompolinsky et al. (2001). Briefly, tuning heterogeneity poses more energy on the high-frequency components of the variance-normalized derivative of mean population response. The more energy on high-frequency components, the more prominent the beneficial effect of TCNC will be. Here, we extend previous work and demonstrate several novel aspects of NC in both neuronal and voxel populations. First, previous theoretical work in neurophysiology primarily focused on estimation tasks (but see (Moreno-Bote et al., 2014)) while the majority of neuroimaging research focused on classification tasks. We compared both tasks in both populations. Second, previous work only analyzed one type of TCNC (i.e., aTCNC in theoretical work) and we systematically compared three types of NC in both populations. Third, we manipulated cTCNC at both neuronal and voxel activity stages to approximate more realistic interaction between neuronal and fMRI responses. These efforts are not only good supplements for existing work in neurophysiology, but also provide a theoretical foundation to understand the effects of NCs in multivariate fMRI data.

### Quantifying information in fMRI data

In this paper, we focused on two routinely used perceptual tasks—the stimulus-estimation task and the stimulus-classification task. Stimulus estimation is equivalent to a very fine-discrimination task as it needs to discriminate the true stimulus value from nearby stimuli in the feature pace. It is primarily determined by Fisher information. Binary classification is more similar to coarse discrimination as it depends on the distance of the representations of two stimuli. In classification tasks, linear discriminability is a better measure than Fisher information (Britten et al., 1992; Butts & Goldman, 2006; Lin et al., 2015; Newsome et al., 1989).

Our work presents a good example to apply this approach in future fMRI research. For example, in a typical binary classification task, we usually measure many trials of population responses towards each of two stimuli. There might be some NCs among voxels but whether they impair or improve population codes is hard to delineate as we have shown that it depends on several factors such as the strength, form of NCs and the pool size. We can use Eq. 18 to tackle this issue. The response difference and covariance matrices can be directly estimated from empirical data. We can therefore use these variables to directly quantify the amount of information. To illustrate the contribution of NCs, we need to compare two conditions—one condition in which estimated response difference, individual response variance, and voxelwise NC are used to calculate the information, the other condition in which the estimated response difference and individual response variance are kept same but the noise correlation matrix is set to diagonal (i.e., all voxelwise NCs are 0). We can conclude that these NCs are detrimental if the removal of NCs elevates the amount of information or vice versa (Averbeck et al., 2006).

### Towards a generative understanding of multivariate fMRI responses

In contrast to the enthusiasm for characterizing generative processes of stimulus-evoked responses in neurophysiology, generative modeling of fMRI data has generally been lacking. Conventional neuroimaging approaches use MVPA to decode information from fMRI data (Haxby et al., 2014; Tong & Pratte, 2012). However, in recent years, people have increasingly realized the limitations of MVPA as a discriminative modeling approach, in which one seeks to estimate the probability *p*(stimulus | response). Rich representational information might be buried by merely examining decoding accuracy (Naselaris & Kay, 2015).

From a probabilistic modeling perspective, understanding the generative computation in the brain is equivalent to deriving the joint probability *p*(response, stimulus), equivalent to *p*(response | stimulus) × *p*(stimulus) according to Bayes’ theorem. Current voxel-encoding modeling approaches seek to characterize the mean of the likelihood term, *p*(response | stimulus), in the sense of characterizing the computations by which a stimulus produces population responses in the brain. However, the full likelihood function *p*(response | stimulus) also requires characterizing the covariance between voxels. Calculating the full likelihood or joint distribution of responses and stimuli can provide important insight into the probabilistic computation in the human brain (van Bergen et al., 2015).

### The nature of noise correlations in fMRI data

Although fMRI can naturally measure the activity of many units in the brain, the investigation of NCs in fMRI data has just begun recently. Exploring this issue in fMRI data is, however, non-trivial and we summarize the related issues as follows.

First, the definition of “noise correlation” in fMRI research is still under debate. The well-accepted definition of “noise correlation” in computational neuroscience is the correlation of trial-by-trial responses between two neurons given the repeated presentation of the same stimulus. This definition emphasizes stimulus-evoked responses. In this paper, we strictly follow this definition and assume voxel responses as trial-by-trial responses estimated from the standard general linear model. This is also called “beta series correlation” in some fMRI literature (Rissman et al., 2004). In contrast, one recent study defined the noise correlation between two voxels as their resting-state functional connectivity or background functional connectivity during a task (Bejjanki et al., 2017). In theory, these definitions deviate from the conventional definition in computational neuroscience and their quantitative relationship remains unclear. Only one recent study suggested that resting-state functional connectivity is highly correlated with the trial-by-trial response correlation at the whole-brain level (Di et al., 2018). Future studies need to examine the relations between resting-, task-based functional connectivity, and trial-by-trial variation of responses at the individual voxel level.

Second, the sources of noise correlations in fMRI data are still unclear. On one hand, the conventional term “noise correlation” itself is somewhat misleading since, as shown in this paper, response variability can contain a substantial amount of stimulus information. In other words, response variability is not purely “noise” and might reflect some critical aspects of how neurons process stimulus or task structure (Bondy et al., 2018). On the other hand, empirically measured voxel response variability also contains non-neural types of noise, such as thermal noise, physiological motion, head motion, etc. These forms of noise are irrelevant to neural information processing but will affect decoding results. Thus, it remains a challenge for future studies to carefully disentangle these different sources of noise correlation.

Third, unlike the established empirical measurements of NC in neurophysiological data, the nature of NC in fMRI data and how it is related to the tuning similarity of voxels still needs to be quantified. Some literature has provided some insight into the relationship between voxel tuning and NCs (Ryu & Lee, 2018; van Bergen & Jehee, 2018). Future studies are needed to investigate this issue in more brain areas, feature dimensions, and tasks.

## MATERIALS AND METHODS

Previous endeavors of brain decoding generally fall into two broad categories: classification of stimuli into discrete categories (Kamitani & Tong, 2005) and estimation of a continuous stimulus variable (Vintch & Gardner, 2014). We thus evaluated the effect of NC in brain decoding in two tasks—a stimulus-classification task and a stimulus-estimation task. We will first introduce the simulation on a neuronal population and then specify the voxel-encoding model used to generate simulated responses of a voxel population.

### Assessment of effects of noise correlations in neuronal populations

#### Neuron-encoding model

The neuron-encoding model assumes 180 orientation-selective neurons whose preferred orientations are equally spaced in [1°, 180°]. Similarly, all orientations throughout the entire paper are angles in degrees within [1°, 180°]. Tuning curves of the neurons can be described as:

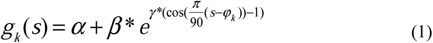

where *g*_*k*_(*s*) is the tuning function of the *k*-th neuron. *s* is the stimulus. φ_*k*_ indicates the preferred orientation of the *k*-th neuron. α is the baseline firing rate, β controls the response range, and γ controls the width of the tuning curve. We set the parameter values α = 1, β = 19 and γ = 2, resulting in a tuning curve with the maximum firing rate at 20 spikes per second. This tuning curve is consistent with previous theoretical work (Ecker et al., 2011) and empirical measurements in the primary visual cortex in primates (Ecker et al., 2010).

Based on this setting, the mean of neuronal population responses given stimulus *s* can be represented by ***G***(*s*)=[*g*_*i*_(*s*)]. However, empirically measured neuronal responses vary trial-by-trial. We posit that the mean of trial-by-trial population responses is ***G***(*s*). We will detail the covariance in the following section.

#### Noise correlation and covariance

We proposed three types of NCs for neuronal data: angular-based tuning compatible noise correlation (aTCNC), curve-based tuning compatible noise correlation (cTCNC) and shuffled noise correlation (SFNC).

**Table 1.** List of symbols

| Symbol | Meaning |
| --- | --- |
| NC | Noise correlation |
| SC | Signal correlation |
| fMRI | Functional magnetic resonance imaging |
| MVPA | Multivariate pattern analysis |
| TCNC | Tuning-compatible noise correlation |
| aTCNC | Angular-based tuning-compatible noise correlation |
| cTCNC | Curve-based tuning-compatible noise correlation |
| SFNC | Shuffled noise correlation |
| $R^{aTCNC}$ | Angular-based tuning-compatible noise correlation matrix |
| $R^{cTCNC}$ | Curve-based tuning-compatible noise correlation matrix |
| $R^{SFNC}$ | Shuffled noise correlation matrix |
| $C_{neuron}$ | Noise correlation coefficient between neurons |
| $C_{vox}$ | Noise correlation coefficient between voxels |
| $C_{homo}$ | voxel tuning heterogeneity coefficient |
| $W$ | Linear weighting matrix from neuronal to voxel responses |

Several theoretical studies assume the NC between a pair of neurons is an exponential function of the angular difference between their preferred orientations, here defined as angular-based tuning compatible noise correlation (aTCNC):

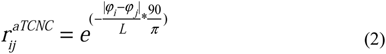

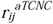 is the NC between the *i*-th and the *j*-th neurons. φ_*i*_ and φ_*j*_ are their preferred orientation. This equation specifies that the NC between two neurons diminishes as their preferred orientations are farther apart. The parameter *L* controls the magnitude of such decay. We denote the correlation matrix as *R*^*a*TCNC^. Here we set *L* = 1 for simplicity. Ecker et al. (2011) has shown that the parametric form of NC and the value of *L* does not qualitatively change the result of the simulation, as long as the generated correlation matrix is positive definite. Note that by this definition aTCNCs are always positive (i.e., range 0~1, also see Fig. 3A).

The second type is the curve-based tuning compatible noise correlation (cTCNC). In this case, the NC between a pair of neurons is proportional to their SC (i.e., correlation of their orientation tuning curves):

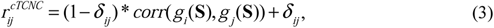

where *δ*_*ij*_ is the Kronecker delta (*δ*_*ij*_=1 if *i* =*j* and *δ*_*ij*_ = 0 otherwise). **S** indicates all possible orientations between [1°, 180°], and 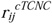 is the NC between the *i*-th and the *j*-th neurons. *g*_*i*_(**S**) and *g*_*j*_(**S**) are their tuning curves (see Eq. 1). We denote *R*^*c*TCNC^ as the correlation matrix. Note that unlike aTCNCs, cTCNCs can be negative (see Fig. 3B). Also, the key difference between cTCNC and aTCNC is that cTCNC does not rely on the functional form of tuning curves. In other words, cTCNC can be computed given irregular tuning curves, whereas aTCNC can be only computed from unimodal tuning curves. This is important for specifications of voxelwise NCs (see below).

In the third case, we shuffled the NCs between all pairs of neurons in *R*^*c*TCNC^ such that the rows and columns are rearranged in the same randomized order but the diagonal of the matrix is kept intact (Fig. 3C). We term this type of NC as shuffled noise correlation (SFNC) since the correlation is no longer necessarily related to the neuronal tuning relations. The correlation matrix of SFNCs is denoted as *R*^*SFNC*^. *R*^*SFNC*^ can serve as a comparison for *R*^*c*TCNC^ since shuffling does not alter the overall distribution of NCs in a neuronal population.

Furthermore, we assumed Poisson noise of spikes such that the response variance of a neuron is equal to the mean activity evoked by a stimulus.

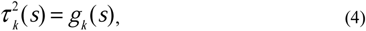

where 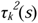 is the response variance of the *k*-th neuron triggered by the stimulus *s*. Note that in this case the response variance is stimulus-dependent. The covariance between neurons *i* and *j* (*q*_*neuron_ij*_ as below) can be expressed as:

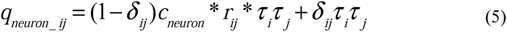

where *c*_*neuron*_ is a parameter that controls the strength of the neuronal NC. *τ*_*i*_ and *τ*_*j*_ are the standard deviation of responses of the two neurons (see Eq. 4), respectively. *δ*_*ij*_ is the Kronecker delta. Given the covariance matrix **Q**_*neuron*_, we can express the population response noise distribution as:

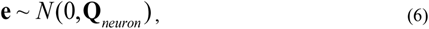

#### Data simulation and multivariate pattern analysis

##### Stimulus-classification task

In the stimulus-classification task, we attempted to determine which of two stimuli were presented, based on the simulated neuronal population responses. We manipulated two independent variables: population size (i.e., the number of neurons) and NC strength (i.e., *c*_*neuron*_ in Eq. 5). We built a linear discriminant using the Matlab function *classify.m*. The linear discriminant assumes that the conditional probability density functions *p*(**b** | *s* = *s*_1_) and *p*(**b** | *s* = *s*_2_) are both normally distributed with the same covariance and estimates the means and covariance from the training data. Here **b** is the vector of a population response in one trial (also see Eq. 7). The classifier was trained on half of the data and tested on the other half.

For the neuronal populations, we attempted to classify two stimuli: *s*_*1*_ = 92°, *s*_*2*_ = 88°. The two stimuli were chosen to control the overall task difficulty (i.e., avoid ceiling and floor effects in classification accuracy). We set six pool size levels (i.e., 10, 20, 50, 100, 200, and 400 neurons) and six NC strength levels (i.e., *c*_*neuron*_ = 0, 0.1, 0.3, 0.5, 0.7, and 0.9). For each combination of a pool size and a *c*_*neuron*_ value and for each form of NC, we performed 100 independent simulations and then averaged classification accuracy values across simulations. To compensate for potential overfitting as the pool size increases, we set the number of trials for each stimulus to be 100 times the pool size. All data were equally divided into two independent parts for training and testing.

##### Stimulus-estimation task

In the stimulus-estimation task, neuronal responses in a trial were simulated for an orientation randomly chosen within [1°, 180°], and then a maximum likelihood estimator (MLE) was used to reconstruct the orientation value. Formally, given a population response pattern **b** in a trial, we attempted to find the stimulus *s* that maximizes the likelihood:

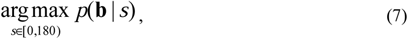

Note that the likelihood function has been introduced above as the neuron-encoding model (see noise distribution in Eqs. 5&6). We numerically evaluated the likelihood of a pattern response **b** for each of 180 integer stimulus orientations (i.e., 1-180°) and chose the orientation that yielded the maximum likelihood value. It is worth noting that, in contrast to classification, the MLE method does not involve any model training, and estimations were directly performed based on the known generative neuron-encoding model. We randomly sampled 1000 stimuli (i.e., 1000 trials) from [1°,180°] for decoding. The same pool size and *c*_*neuron*_ settings as in the stimulus-classification task were used. For each combination of a pool size and a *c*_*neuron*_ value, we calculated the mean circular squared errors (*MSE*_*circ*_) across all trials between the estimated stimuli 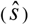 and the true stimuli (*s*) across all trials:

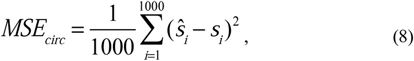

where 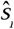 is the estimated stimulus and *s*_*i*_ is the true stimulus in the *i*-th trial. We took the inverse of the *MSE*_*circ*_ as the estimation efficiency (see Fig. 4&5). A higher estimation efficiency value indicates a more accurate estimation.

### Assessment of effects of noise correlations in voxel populations

#### Voxel-encoding model

The voxel-encoding model uses the same pool of orientation-selective neurons (i.e., 180 neurons with tuning curves defined in Eq. 1) as in the neuron-encoding model. We further assume that the response of a voxel is the linear combination of all neurons in the neuronal population:

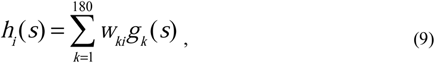

where *h*_*i*_(*s*) is the tuning function of the *i*-th voxel. *w*_*ki*_ is the connection weight between the *k*-th neuron to the *i*-th voxel. We sampled *w*_*ki*_ from a uniform distribution:

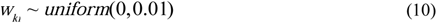

This range was used so that generated fMRI responses typically range between 0 and 10 (and can be viewed as approximating units of percent BOLD change). This is also consistent with the range of empirically measured fMRI responses in most studies.

The mean of voxel population response given stimulus *s* can be represented by ***H***(*s*)=[*h*_*i*_(*s*)]. To express the trial-by-trial variation of voxel responses, we specify:

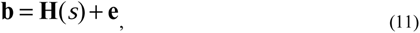

Here, **b** represents the observed response across voxels on a trial (as might be obtained from a general linear model applied to fMRI data) and **e** represents the multivariate normal noise distribution:

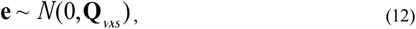

where **Q**_*vxs*_ is the covariance matrix between voxels, which will be detailed in the following section.

#### Noise correlation and covariance

We evaluate two types of NCs for simulated fMRI data: cTCNC and SFNC. Note that we cannot evaluate aTCNC for voxel populations because voxel tuning curves here are irregular and not unimodal (see Fig. 8C).

In the first case, we defined cTCNC using a similar method as Eq. 3:

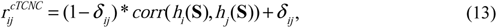

where 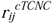 is the NC between voxels *i* and *j*. Note that the cTCNC here is based on the tuning curves of two voxels (i.e., *h*_i_(**S**) and *h*_j_(**S**)), not two neurons. *δ*_*ij*_ is the Kronecker delta.

In the second case, SFNCs were generated using a similar method as in the neuron-encoding model—shuffling the rows and columns in *R*^*c*TCNC^, which is obtained in Eq. 13.

We assume the response variances for different voxels (e.g, 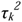 for the *k*-th voxel) follow a Gamma distribution:

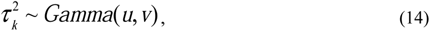

where *u* = 9, *v* = 0.33 are the scale and the shape parameters corresponding to a Gamma distribution with mean = 3 and variance = 1. Given the response variance of individual voxels and the NC between them, we can calculate the covariance between the *i*-th and the *j*-th voxels as:

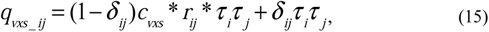

where *c*_*vxs*_ is the parameter that controls the strength of the voxelwise NCs. *τ*_*i*_ and *τ*_*j*_ are the standard deviation of responses of the two voxels (from Eq. 14), respectively. *δ*_*ij*_ is the Kronecker delta. Given the covariance matrix **Q**_vxs_, we can finally generate voxel population responses using Eqs. 11&12. Note that Eq. 14 describes the variability of the response variance across voxels. The distribution of voxel population responses still follows a multivariate Gaussian distribution (Eq. 12).

#### Data simulation and multivariate pattern analysis

##### Stimulus-classification task

In the voxel-encoding model, we reduced the task difficulty and set the two stimuli as *s*_*1*_ = 80°, *s*_*2*_ = 100°. The motivation for changing task difficulty is to compensate for the higher noise level in voxel responses and avoid ceiling or floor effects in classification. We set six pool size levels (i.e., 10, 20, 50, 100, 200, and 500 voxels) and eight NC strength levels (i.e., *c*_*vxs*_ = 0, 0.01, 0.03, 0.1, 0.3, 0.5, 0.7, and 0.9). 10 independent simulations were performed. In each simulation, we assessed the classification accuracy for each combination of a pool size and a *c*_*vxs*_ value. Since the voxel tuning curves are determined by the linear weighting matrix **W**, in each simulation, we generated a new **W** for a given pool size. This ensures that we generated a new set of voxels in every simulation such that our conclusion is not biased by a particular choice of **W**. The **W** was kept constant across different *c*_*vxs*_ values such that classification accuracy values are directly comparable across different *c*_*vxs*_ values. For each stimulus, we simulated 1000, 1000, 1000, 1000, 2000, and 5000 trials for the corresponding pool sizes, respectively. We increased the number of trials for large pool sizes to avoid overfitting.

##### Stimulus-estimation task

In the stimulus-estimation task, we used the same pool size and NC strength settings as in the neuron-encoding model. We also performed 10 independent simulations and generated simulated responses to 1000 stimuli between [1°, 180°] in each simulation. Similar to above, for each simulation and each pool size, we recreated a linear weight **W** to create a new set of voxels, and kept the same **W** across *c*_*vxs*_ values. Similar to neuronal populations, the inverse of circular mean square error (Eq. 8) was calculated to indicate the estimation efficiency. The estimation efficiency values were averaged across the 10 simulations.

### Simultaneously manipulating neuronal and voxelwise noise correlations

In previous simulations, we either only manipulated the neuronal NCs in the neuron-encoding model or the voxelwise NCs in the voxel-encoding model. However, in realistic fMRI responses, voxel responses will inherit NCs from the neural level and will also include other sources of NC (such as head motion). To examine the interaction between neuron-level and voxel-level NCs on decoding accuracy, we simultaneously manipulated both neuronal and voxelwise NCs in the voxel-encoding model (Fig. 6). In this simulation, we kept the same settings as the simulations above in the voxel-encoding model except for the following changes. First, we fixed the pool size to 200 voxels and manipulated the cTCNCs at the neuron level. We set eight cTCNC strength levels at the neural level (same as in the neuron-encoding model). Second, in every trial of classification or estimation, we first generated a neuronal population response (i.e., responses for 180 neurons). Note that this generation takes into account neuron-level NCs. We then linearly transformed the neuronal population response into a voxel population response using the linear weighting matrix **W** (10 different **W** for 10 independent simulations), which yields the mean of the voxel population response. Finally, to generate the voxel population response observed on a given trial, we added voxel-level cTCNCs to the mean voxel population response, as we did in previous simulations.

### Differences between the neuron- and the voxel-encoding models

In this section, we summarize three key differences between the two encoding models proposed above. The biggest difference is that decoding is performed directly on simulated neuronal responses in the neuron-encoding model, but is performed on simulated voxel responses, which are linear combinations of the underlying neuronal responses, in the voxel-encoding model. Second, the NCs we manipulate are between neurons in the neuron-encoding model. In the voxel-encoding model, we either only manipulate the NCs at the voxel level (Fig. 5) or the NCs in both neuron and voxel stages (Fig. 6). In empirical fMRI studies, neuronal NCs are inaccessible; thus, the former case is more pertinent to realistic fMRI data analysis while the latter case provides theoretical insights. Third, we assume Poisson-like response variance for individual neurons in the neuron-encoding model, which is consistent with the previous theoretical work and empirical findings (Kohn et al., 2016). In this regime, the magnitude of response variance of individual neurons is stimulus-dependent. In the voxel-encoding model, we assume stimulus-independent additive Gaussian noise for voxels, consistent with one recent computational study (van Bergen & Jehee, 2018).

### Information-theoretic analyses

We calculated Fisher information in the stimulus-estimation task as it is one of the standard methods to quantify information in computational neuroscience (Brunel & Nadal, 1998). Specifically, we used linear Fisher information, which is a linear approximation of Fisher information, and can be expressed as:

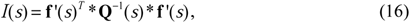

where **f***’*(*s*) is the derivative of the mean population responses with respect to stimulus *s* and **Q (**i.e., **Q**_*neuron*_ or **Q**_*vxs*_) is the covariance matrix given stimulus *s*. Note that linear Fisher information can be calculated from both simulated neuron- and voxel-encoding models as long as the tuning curves and the covariance matrix are known. Notably, in neuronal data, complete Fisher information is stimulus-dependent because of the assumed Poisson noise distribution and the covariance matrix **Q** varies across stimuli. Thus, linear Fisher information here is not identical to complete Fisher information but previous studies have shown it is a good approximation (Moreno-Bote et al., 2014; Kohn et al., 2016). In the simulated voxel data, we assumed additive Gaussian noise and thus the covariance matrix **Q** is identical for all orientations (i.e., stimulus-invariant) and thus linear Fisher information is equivalent to complete Fisher information. In this paper, we simply denote both as “information”. We computed the averaged linear Fisher information for all 180 discrete orientations:

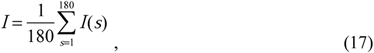

where *s* is the stimulus. Note that theoretically linear Fisher information above only applies to the stimulus-estimation task or a fine-discrimination task. For a general classification task (i.e., classify two stimuli *s*_1_ and *s*_2_), especially for a coarse discrimination task, Eq.16 can be rewritten as the discrete format:

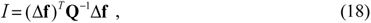

where Δ**f** = **f**(*s*_1_) − **f**(*s*_2_) is the population response difference for two stimuli and 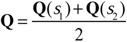 is the covariance matrix. This metric by definition is not Fisher information and typically called “linear discriminability” (Lin et al., 2015). To avoid confusion in terminologies, we denoted both metrics as “information” in this paper as they indicate the quality of population codes in the two tasks respectively. In the main text, we only show the information in the stimulus-estimation task (Fig. 6).

### Varying voxel tuning heterogeneity

To illustrate the effect of tuning heterogeneity, we performed an additional analysis on the voxel-encoding model (Fig. 8). In this analysis, we calculated the amount of information in the stimulus-estimation task after making the following modifications. First, we fixed the voxel pool size to 500. Second, we introduced the heterogeneity coefficient (*c*_*homo*_) that controls the voxel tuning heterogeneity. The key to manipulating heterogeneity is to adjust the linear weighting **W** from neuronal to voxel responses. For each voxel, we first randomly selected one neuron from all 180 neurons and assigned *c*_*homo*_ as the linear weight for this neuron. The weights for other neurons were then assigned by random numbers between 0~1 scaled by (1- *c*_*homo*_) (i.e., (1- *c*_*homo*_)*rand in Matlab). For example, if *c*_*homo*_ = 1, the voxel tuning curve is homogeneous and identical to the neuronal tuning curve chosen in the first step; if *c*_*homo*_ = 0, the voxel tuning curves are heterogeneous as it is the linear combination of all other neurons with random weights (see Fig. 8C). Third, one might speculate that differences in results across neuronal and voxel simulations might due to the absolute response range. In the neuron-encoding models, the response range of neuronal tuning curves is between [1, 20] spikes per second whereas the voxel tuning curves are smaller than 10. To control this absolute difference in the response ranges, we normalized the range of voxel tuning curves to [1, 20] (see Eq. 1, also see scaled voxel tuning curves in Fig. 9C compared to Fig. 8C). Note that in this case voxel response amplitude is larger than that in the previous voxel simulation (<10). Larger response amplitudes will result in overall higher information, we thus also scaled the voxel variance 40 times (i.e., the mean of Gamma distribution in Eq. 14) to keep the comparable signal-to-noise levels in voxel responses.

## Code availability

Relevant code for the simulations have been available at https://github.com/ruyuanzhang/noisecorrelation

## ACKNOWLEDGMENTS

The work was supported by NIH Grants P41 EB015894, P30 NS076408, S10 RR026783, S10 OD017974-01, NSF NeuroNex Award DBI-1707398, the Gatsby Charitable Foundation, and the W.M. Keck Foundation.

## AUTHOR CONTRIBUTIONS

R-Y. Z. conceived, designed and performed research; R-Y. Z. wrote the draft of the paper. R-Y. Z., X-X. W., and K.K. edited the paper.

## REFERENCES

Averbeck, B. B., Latham, P. E., & Pouget, A. (2006). Neural correlations, population coding and computation. Nat Rev Neurosci 7:358–366. https://doi.org/10.1038/nrn1888

Averbeck, B. B., & Lee, D. (2003). Neural noise and movement-related codes in the macaque supplementary motor area. J Neurosci 23:7630–7641.

Bair, W., Zohary, E., & Newsome, W. T. (2001). Correlated firing in macaque visual area MT: time scales and relationship to behavior. J Neurosci 21:1676–1697. https://doi.org/10.1523/JNEUROSCI.21-05-01676.2001

Bejjanki, V. R., da Silveira, R. A., Cohen, J. D., & Turk-Browne, N. B. (2017). Noise correlations in the human brain and their impact on pattern classification. PLoS Comput Biol 13:e1005674. https://doi.org/10.1371/journal.pcbi.1005674

Bondy, A. G., Haefner, R. M., & Cumming, B. G. (2018). Feedback determines the structure of correlated variability in primary visual cortex. Nat Neurosci 21:598–606. https://doi.org/10.1038/s41593-018-0089-1

Britten, K. H., Shadlen, M. N., Newsome, W. T., & Movshon, J. A. (1992). The analysis of visual motion: a comparison of neuronal and psychophysical performance. J Neurosci 12:4745–4765. https://doi.org/10.1523/JNEUROSCI.12-12-04745.1992

Brunel, N., & Nadal, J. P. (1998). Mutual information, Fisher information, and population coding. Neural Comput 10:1731–1757. https://doi.org/10.1162/089976698300017115

Butts, D. A., & Goldman, M. S. (2006). Tuning curves, neuronal variability, and sensory coding. PLoS Biol 4:e92. https://doi.org/10.1371/journal.pbio.0040092

Cohen, M. R., & Kohn, A. (2011). Measuring and interpreting neuronal correlations. Nat Neurosci 14:811–819. https://doi.org/10.1038/nn.2842

Constantinidis, C., & Goldman-Rakic, P. S. (2002). Correlated discharges among putative pyramidal neurons and interneurons in the primate prefrontal cortex. J Neurophysiol 88:3487–3497. https://doi.org/10.1152/jn.00188.2002

Cox, D. D., & Savoy, R. L. (2003). Functional magnetic resonance imaging (fMRI)“brain reading”: detecting and classifying distributed patterns of fMRI activity in human visual cortex. NeuroImage 19:261–270.

Di, X., Zhang, Z., & bioRxiv, B.-B. B. (2018). Psychophysiological interaction and beta series correlation for task modulated connectivity: modeling considerations and their relationships. bioRxiv:322073. https://doi.org/10.1101/322073

Ecker, A. S., Berens, P., Keliris, G. A., Bethge, M., Logothetis, N. K., & Tolias, A. S. (2010). Decorrelated neuronal firing in cortical microcircuits. science 327:584–587. https://doi.org/10.1126/science.1179867

Ecker, A. S., Berens, P., Tolias, A. S., & Bethge, M. (2011). The effect of noise correlations in populations of diversely tuned neurons. J Neurosci 31:14272–14283. https://doi.org/10.1523/JNEUROSCI.2539-11.2011

Graf, A. B., Kohn, A., Jazayeri, M., & Movshon, J. A. (2011). Decoding the activity of neuronal populations in macaque primary visual cortex. Nat Neurosci 14:239–245. https://doi.org/10.1038/nn.2733

Gutnisky, D. A., & Dragoi, V. (2008). Adaptive coding of visual information in neural populations. Nature 452:220–224. https://doi.org/10.1038/nature06563

Haefner, R. M., Gerwinn, S., Macke, J. H., & Bethge, M. (2013). Inferring decoding strategies from choice probabilities in the presence of correlated variability. Nat Neurosci 16:235–242. https://doi.org/10.1038/nn.3309

Haxby, J. V., Connolly, A. C., & Guntupalli, J. S. (2014). Decoding neural representational spaces using multivariate pattern analysis. Annu Rev Neurosci 37:435–456. https://doi.org/10.1146/annurev-neuro-062012-170325

Huang, X., & Lisberger, S. G. (2009). Noise correlations in cortical area MT and their potential impact on trial-by-trial variation in the direction and speed of smooth-pursuit eye movements. J Neurophysiol 101:3012–3030. https://doi.org/10.1152/jn.00010.2009

Jermakowicz, W. J., Chen, X., Khaytin, I., Bonds, A. B., & Casagrande, V. A. (2009). Relationship between spontaneous and evoked spike-time correlations in primate visual cortex. J Neurophysiol 101:2279–2289. https://doi.org/10.1152/jn.91207.2008

Kamitani, Y., & Tong, F. (2005). Decoding the visual and subjective contents of the human brain. Nat Neurosci 8:679–685. https://doi.org/10.1038/nn1444

Kohn, A., Coen-Cagli, R., Kanitscheider, I., & Pouget, A. (2016). Correlations and Neuronal Population Information. Annu Rev Neurosci 39:237–256. https://doi.org/10.1146/annurev-neuro-070815-013851

Lee, D., Port, N. L., Kruse, W., & Georgopoulos, A. P. (1998). Variability and correlated noise in the discharge of neurons in motor and parietal areas of the primate cortex. J Neurosci 18:1161–1170.

Lin, I. C., Okun, M., Carandini, M., & Harris, K. D. (2015). The Nature of Shared Cortical Variability. Neuron 87:644–656. https://doi.org/10.1016/j.neuron.2015.06.035

Ma, W. J., & Jazayeri, M. (2014). Neural coding of uncertainty and probability. Annu Rev Neurosci 37:205–220. https://doi.org/10.1146/annurev-neuro-071013-014017

Moreno-Bote, R., Beck, J., Kanitscheider, I., Pitkow, X., Latham, P., & Pouget, A. (2014). Information-limiting correlations. Nat Neurosci 17:1410–1417. https://doi.org/10.1038/nn.3807

Naselaris, T., & Kay, K. N. (2015). Resolving Ambiguities of MVPA Using Explicit Models of Representation. Trends Cogn Sci 19:551–554. https://doi.org/10.1016/j.tics.2015.07.005

Naselaris, T., Kay, K. N., Nishimoto, S., & Gallant, J. L. (2011). Encoding and decoding in fMRI. NeuroImage 56:400–410. https://doi.org/10.1016/j.neuroimage.2010.07.073

Newsome, W. T., Britten, K. H., & Movshon, J. A. (1989). Neuronal correlates of a perceptual decision. Nature 341:52–54. https://doi.org/10.1038/341052a0

Rissman, J., Gazzaley, A., & D’Esposito, M. (2004). Measuring functional connectivity during distinct stages of a cognitive task. NeuroImage 23:752–763. https://doi.org/10.1016/j.neuroimage.2004.06.035

Ryu, J., & Lee, S. H. (2018). Stimulus-Tuned Structure of Correlated fMRI Activity in Human Visual Cortex. Cereb Cortex 28:693–712. https://doi.org/10.1093/cercor/bhw411

Seung, H. S., & Sompolinsky, H. (1993). Simple models for reading neuronal population codes. Proc Natl Acad Sci U S A 90:10749–10753. https://doi.org/10.1073/pnas.90.22.10749

Shamir, M., & Sompolinsky, H. (2006). Implications of neuronal diversity on population coding. Neural Comput 18:1951–1986. https://doi.org/10.1162/neco.2006.18.8.1951

Smith, M. A., & Kohn, A. (2008). Spatial and temporal scales of neuronal correlation in primary visual cortex. J Neurosci 28:12591–12603. https://doi.org/10.1523/JNEUROSCI.2929-08.2008

Sompolinsky, H., Yoon, H., Kang, K., & Shamir, M. (2001). Population coding in neuronal systems with correlated noise. Phys Rev E Stat Nonlin Soft Matter Phys 64:051904. https://doi.org/10.1103/PhysRevE.64.051904

Tong, F., & Pratte, M. S. (2012). Decoding patterns of human brain activity. Annu Rev Psychol 63:483–509. https://doi.org/10.1146/annurev-psych-120710-100412

van Bergen, R. S., & Jehee, J. F. M. (2018). Modeling correlated noise is necessary to decode uncertainty. NeuroImage 180:78–87. https://doi.org/10.1016/j.neuroimage.2017.08.015

van Bergen, R. S., Ma, W. J., Pratte, M. S., & Jehee, J. F. (2015). Sensory uncertainty decoded from visual cortex predicts behavior. Nat Neurosci 18:1728–1730. https://doi.org/10.1038/nn.4150

Vintch, B., & Gardner, J. L. (2014). Cortical correlates of human motion perception biases. J Neurosci 34:2592–2604. https://doi.org/10.1523/JNEUROSCI.2809-13.2014

Wilke, S. D., & Eurich, C. W. (2002). Representational accuracy of stochastic neural populations. Neural Comput 14:155–189. https://doi.org/10.1162/089976602753284482

Zohary, E., Shadlen, M. N., & Newsome, W. T. (1994). Correlated neuronal discharge rate and its implications for psychophysical performance. Nature 370:140–143. https://doi.org/10.1038/370140a0

